# Innovative Strategies for Enhancing Signal-to-Noise Ratio in Single-molecule Imaging: The Influence of Molecular Motion

**DOI:** 10.1101/2024.10.13.618074

**Authors:** Gayun Bu, Gyeongho Kang, Seongyu Park, Jae-Hyung Jeon, Jong-Bong Lee

## Abstract

Single-molecule fluorescence imaging has extensively revealed the dynamic and structural characteristics of biomolecules. However, its application is limited by the upper concentration of fluorophore-tagged biomolecules, which is in the sub-ten nanomolar range. We found that the signal-to-noise ratio (SNR) in single-molecule fluorescence imaging is strongly influenced by the size of fluorophore-labeled molecules in solution. Our computational simulations suggest that the faster diffusion of background fluorophores can enhance the SNR of target molecules. Moreover, we identified that the molecular motion through fluid flow can improve SNR. This study provides a novel perspective by emphasizing the importance of molecular motion in SNR and propose a rapid barrier-free method to increase the upper concentration limit in single-molecule imaging.

## Introduction

Visualizing of a single fluorophore in single-molecule fluorescence imaging requires a high signal-to-noise ratio (SNR) to detect individual fluorophore-tagged molecules [1]. To achieve this, total internal reflection fluorescence microscopy (TIRFM) is commonly used in single-molecule experiments, as it provides the smallest illumination volume among wide-field microscopy techniques [2]. However, the feasible concentration of molecules during detection is limited to the tens nanomolar range [3, 4]. Consequently, most single-molecule studies are performed below this upper concentration limit to study the dynamic features of biological events. Alternatively, molecules from the solution are washed out after a certain amount of reaction has occurred.

Many biological systems interact with their binding partners with affinities ranging from tens of nanomolar to micromolar or even millimolar[5]. Investigating the real-time kinetics of these molecules requires their presence in solution, but concentration limitations often make detection challenging. Various approaches have been developed to address these limitations. For instance, using photoactivatable dyes [3, 6, 7] or nanostructure imaging chambers [8-10] has been proposed and successfully implemented. However, these methods necessitate the use of special fluorophores, illumination techniques, or specialized equipment.

In this study, we found that an immobilized fluorophore-tagged molecule exhibited a higher SNR when smaller fluorophore-labeled molecules were present in solution compared to larger ones. Our computer-aid simulation showed that the faster rate of diffusion of the free molecules in solution enhances the SNR of surface-bounded molecules. This is because smaller fluorophore-labeled molecules reduce the standard deviation of the background fluorescent signals caused by their movement in the solution. We demonstrated that the hydrodynamic flow can effectively improve SNR, offering a method adaptable to any single-molecule technique.

## Materials and methods

### Preparation of single fluorophore molecules

Oligonucleotides were purchased from Bioneer (Daejeon, Korea), Bionics (Seoul, Korea), or Integrated DNA Technologies (Coralville, IA, USA) to prepare double stranded DNA (dsDNA) of various nucleotide lengths by annealing 5’-Cy5 conjugated oligonucleotides with complementary non-labeled oligos of 29 nt or 92 nt lengths. Additionally, a 29 nt long Cy5 oligo was employed to produce a 1.3 kbp dsDNA via PCR from a modified pQE-UBE1 plasmid (Addgene, Watertown, MA, USA). The PCR product was purified through ethanol precipitation, and the remaining primers were filtered through an Amicon Ultra Centrifugal filter (Merck, Darmstadt, Germany). For the preparation of a fluorophore-tagged ubiquitin, 6xHis-TEV cleavage sequence-Cys(−1)-ubiquitin was expressed in *E. coli* (NEB, Ipswich, MA, USA, C3037) and then purified using Ni-NTA resin (Cytiva, Marlborough, MA, USA). The purified ubiquitin was labeled with Cy5-maleimide (Cytiva or Lumiprobe, Wan Chai, Hong Kong) through Cys(−1). The 6xHis tag was cleaved by uTEV3 (Addgene, Watertown, MA, USA, #135464), leaving Gly-^Cy5^Cys-ubiquitin. The ubiquitin concentration reported in this study indicates the concentration of Cy5 on ubiquitin, which was measured with a Nanophotometer (Implen), as the degree of labeling was not substantial.

### Single-molecule fluorescence imaging

Total internal reflection fluorescence microscopy (TIRFM) was used for visualizing a single fluorophore. Fluorophores were excited by an evanescent wave from a 638 nm laser (Cobolt MLD 180mW), then the emitted signals were collected through an objective lens (Olympus 60x, NA 1.2, water) with a 1.6x magnifier of an inverted microscope (Olympus IX-71). The excitation beam was excluded by an emission filter (Semrock, 676/37) and filtered signals were detected by an EMCCD (C9100-13, Hamamatsu) with an exposure time of 100 ms using MetaMorph 7.6 (Molecular Devices) software.

The flow chamber used for TIRFM imaging was passivated with 5k Polyethylene glycol (PEG) and PEG-biotin (Laysan Bio), as described in the previous study[11]. Recombinant Albumin (NEB), streptavidin (Thermo Fisher Scientific), and 40 nt oligos with 5’-Cy5 and 3’-biotin were sequentially incubated in the flow chamber and then washed free molecules in the solution with blocking buffer (20 mM Tris pH 8.0, 2 mM EDTA, 50 mM NaCl, 0.0025% Tween-20). An image of the signal molecules without background molecules (0 nM) was captured in imaging buffer [25 mM Tris pH 8.0, 100 mM NaCl, 2 mM Trolox, 5 mM PCA (Sigma-Aldrich), 0.003 U rPCO (Oriental Yeast)], followed by the injection and imaging of the indicated concentration of background molecules in the same imaging buffer at the same position as 0 nM. For the fluid flow experiment, a syringe pump (Harvard Apparatus) was used to control the flow rate. All processes were performed at room temperature (22 °C).

### Analysis of experimental single fluorescence images

The use of 20 nM fluorophore-labeled molecules in this experiment led to a poor SNR due to substantial background signals from free fluorophore-labeled molecules in solution. To enhance the surface-bound fluorescence signals in the flow chamber, the images were processed by averaging N frames using the TrackMate plugin in ImageJ[12]. Each frame 1 to N was included in the first frame of the averaged image, frame 2 to N+1 was included in the second averaged frame, and so forth. We selected fluorescent signals that remained unbleached within the 10-frame window during the averaging process.

Based on the positions of the identified fluorescent signals, the spatial SNR (SSNR) was directly calculated from the original images. For the temporal SNR (TSNR), the fluorescent signals were similarly analyzed using a moving average image with a 5-frame window size. To obtain the fluorescence and background intensities of the same fluorophore, the on-state and off-state were identified at the specific position. For TSNR calculations, the intensity profile of the switched-on fluorescence served as the signal, while the off-state fluorescence was treated as the background. The TSNR was calculated at these specific locations from the original image. For SSNR calculations, a region of 19 × 19 pixels was extracted around each fluorescent signal. The analysis was restricted to include only a single fluorescent molecule in this region, as additional fluorescent molecules nearby would compromise the SNR. Similarly, for TSNR, a 7×7 pixel region was isolated, ensuring that only one molecule was analyzed in both cases.

### Simulation of freely diffusing fluorescent molecules and calculation of SNR

To address experimental observations, particles in the flow chamber were simulated to undergo 3D thermal diffusion as described by the Stokes-Einstein relation. Initially, paticle positions were randomly assigned, with subsequent positions calculated every 10 μs (Δ*t*). The next position was determined by a multivariable normal distribution with a covariance value based on the diffusion coefficient *D* for each axis (covariance = 2*Ddt*, mean = 0). Periodic boundary conditions were applied along the x- and y-axes, while the z-axis employed a reflecting boundary condition. The total number of simulated trajectories was proportional to the concentration of background molecules, with each trajectory generated independently. For the fluid flow simulation, a linear velocity along the x-axis was applied to each diffusion particle.

To simulate fluorescence emission, a Gaussian profile with a width of 0.25 μm was applied to each point along the particle trajectories. The fluorescence intensity decayed exponentially along the z-axis to account the evanescent field of the TIRFM excitation beam. Emissions were integrated every 100 ms, mimicking the exposure time of the EMCCD. The resulting total intensities were summed over a 48 × 48 pixel grid, with each pixel corresponding to 0.167 μm × 0.167 μm, matching the dimensions of the experimental setup. For SNR calculations, three or nine signal molecules were placed on the surface (z = 0). To model fluid flow, a constant linear velocity along the x-axis was added to trajectory. The experimental flow chamber, with a width of 3 mm and a height of 100 μm, corresponded to a particle speed 0.056 m/s at an applied flow rate of 1 ml/min.

## Results

### Effect of molecular size on SNR in single-molecule imaging

In our experiments, we discovered that individual fluorophore-labeled ubiquitin can be detected even at its high concentrations in single-molecule imaging using TIRF microscopy. This challenges the conventional understanding of concentration limits in single-molecule imaging. Ubiquitin, a small protein with a molecular weight of 8.9 kDa and a diameter of approximately 3 nm (PDB:1UBI) [13], led us to hypothesize that molecular size could influence the SNR in imaging.

To test this, we developed an SNR measurement procedure by analyzing identical signal molecules against background molecules of varying sizes (Fig. 1A). We defined two types of SNR: spatial SNR (SSNR) and temporal SNR (TSNR) [11, 14]. SSNR evaluates the detectability of a particle in a single frame, which is determined by the intensity and variability of the background. To reduce bias, exceptionally bright or dim fluorescent signals were excluded We acquired images of signal molecules without background interference, profiling histograms of all signal fluorophores. Only fluorophores at least 80% of their intensities within one standard deviation of the Gaussian distribution were included in the analysis. Once background molecules were introduced into the flow chamber, the SSNR was calculated based on the positions of the selected fluorophores. TSNR measures the accuracy of intensity measurements over time. For this, we collected 20 frames of “on” and “off” signals at the same positions as detailed in the methods section.

**Fig. 1.**
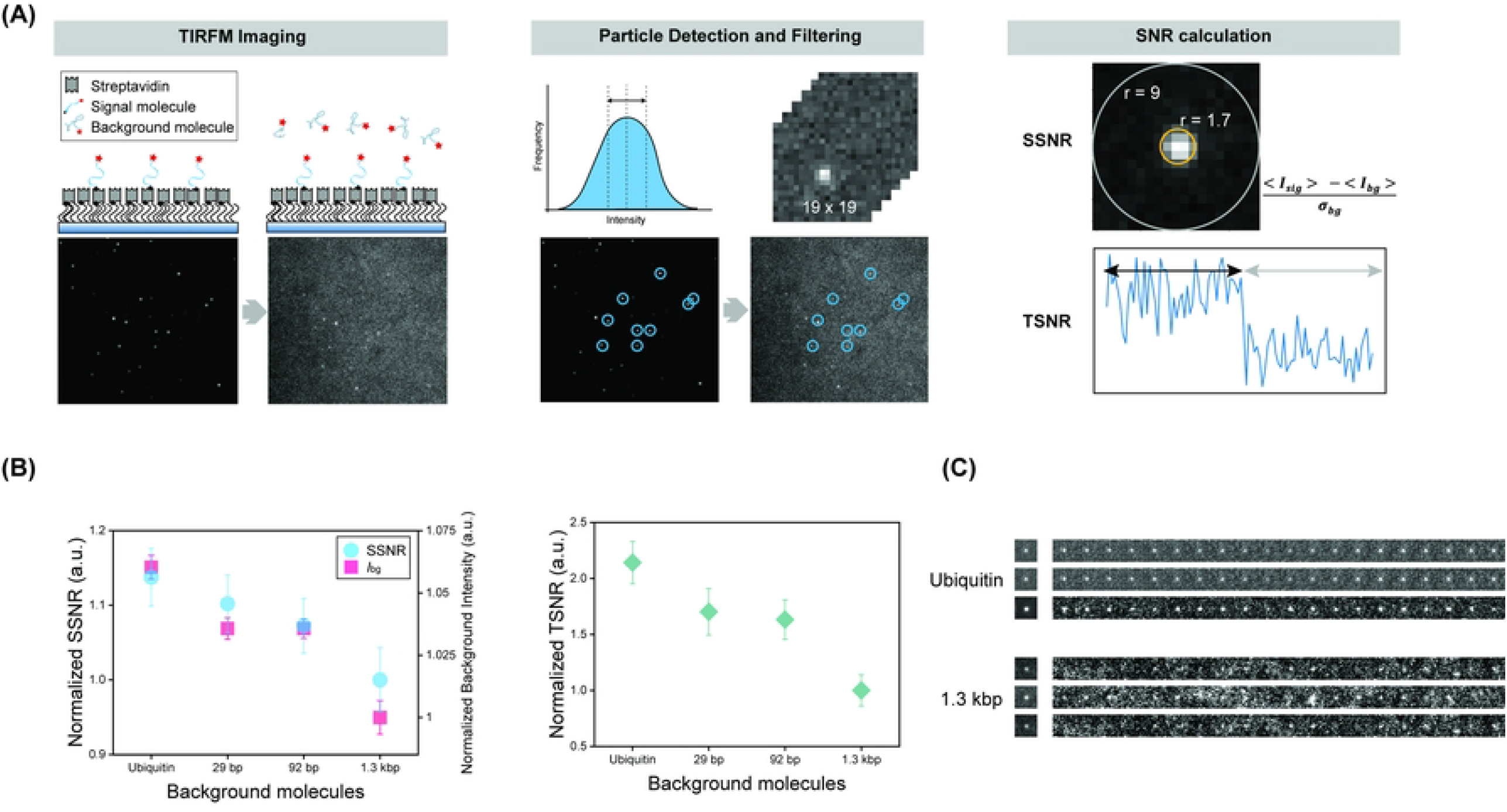
Single-molecule imaging of fluorophore-labeled molecules with different hydrodynamic radii. (A) Schematic representation of the SNR analysis procedure for molecules with varying hydrodynamic radii. (B) (Left) Normalized SSNR for molecules with various hydrodynamic radii and background intensities. N_ubiquitin_ = 98, N_29 bp_ = 118, N_92 bp_ = 112, N_1.3 kbp_ = 103. SSNR for each frame was averaged over all molecules. Error bars indicate standard error. (Right) Normalized TSNR values: N_ubiquitin_ = N_29 bp_ = N_92 bp_ = N_1.3 kbp_ = 11. Since the absolute SNR values varied across different experimental conditions, all SNR values were normalized to those of the 1.3 kbp molecules. (C) Representative images of signals detected in 20 nM ubiquitin and 1.3 kbp dsDNA used as background molecules in solution. The left image is the average of 20 stacks, while the right displays the montage.

SNR was mathematically defined as 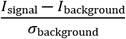, where *I*_signal_ is the average intensity of the signal area, *I*_background_ is the average intensity of the background, and σ_background_ is the standard deviation of the background. For SSNR, the signal area was defined as pixels where *r*^2^ ≤ 3, with *r* being the distance from the center. The background area was defined as 3 < *r*^2^ ≤ 81. For TSNR, we calculated the average intensity from a 7 × 7 pixel grid. The signal area corresponded to frames where fluorophores were “on,” and the background area corresponded to frames where the fluorophores were “off” (Fig. 1A).

It is well established that higher concentrations of background fluorophores, which increase the image’s background intensity, generally lead to lower SNR. However, our findings revealed that smaller fluorophore-labeled molecules, such as ubiquitin, exhibit higher SSNR and TSNR values compared to larger molecules, even at higher concentrations (Fig.1B and C). This trend was consistent across various molecules, including both DNA and proteins.

### Effect of diffusion coefficient on SNR

To explain the improved SNR observed in smaller molecules, we performed a simple computer-aid simulation. In this simulation, molecules undergoing random diffusion were assigned fluorescence emissions with intensities based on an exponential decay as a function of the z-distance from the surface, reflecting the evanescent field generated by the excitation beam in TIRFM (Fig. 2A). According to the Stokes-Einstein relation, 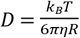, where *D* is the diffusion coefficient, *k*_*B*_ is the Boltzmann constant, *T* is temperature, *η* is viscosity, *R* is the hydrodynamic radius of the molecules. Consequently, molecules with smaller hydrodynamic radii diffuse over a larger area, while larger molecules, with slower diffusion remain more confined.

**Fig. 2.**
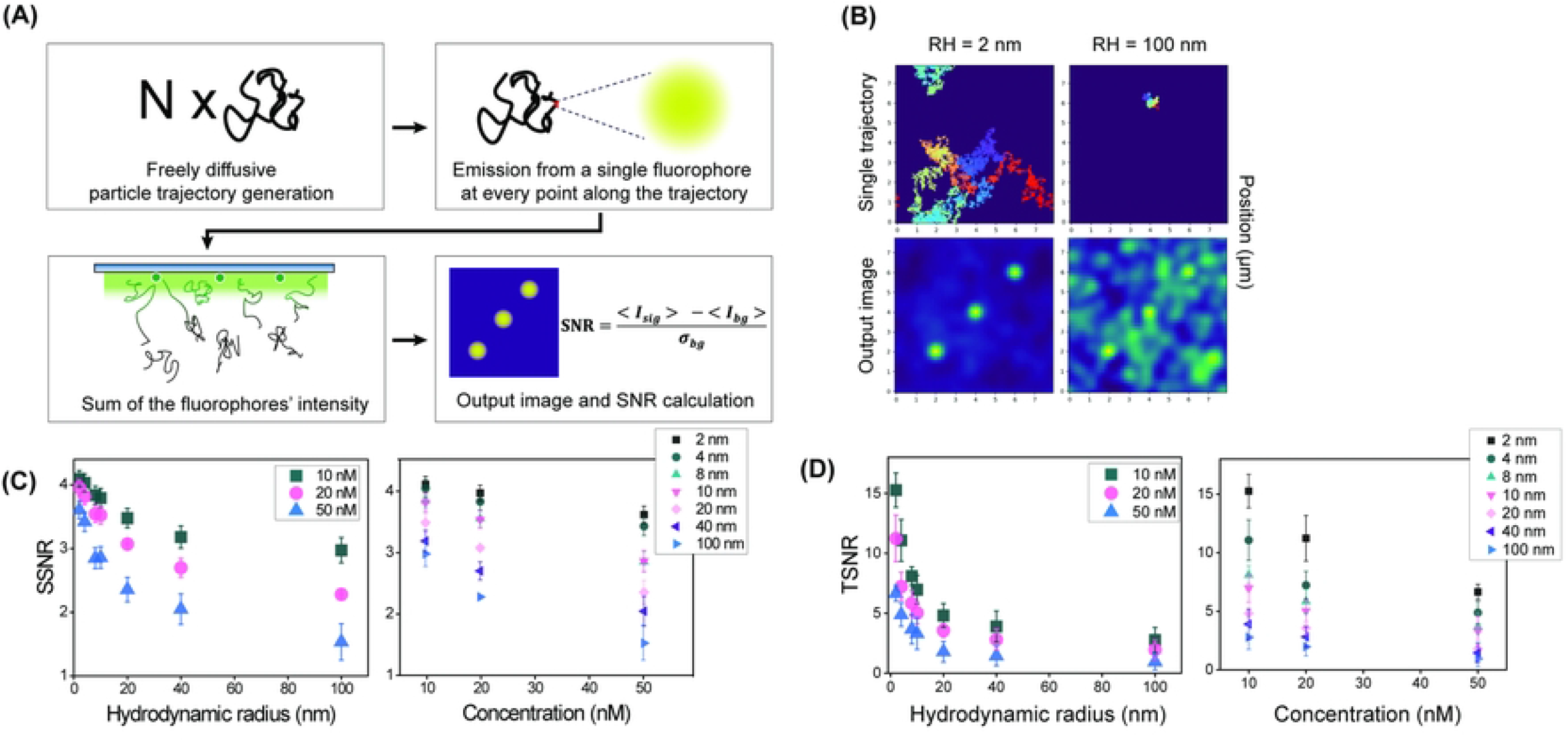
SNR simulation of freely diffusing molecules in different hydrodynamic radii. (A) Schematic for 3D diffusion of molecules and the SNR calculation from output images. (B) Representative single trajectories for background molecules with hydrodynamic radii of 2 nm and 100 nm. Result images were generated as 20 nM concentration. (C), (D) SSNR and TSNR decreased with increasing hydrodynamic radius (left) and concentration (right). For SSNR calculations, 9 molecules were analyzed per frame. For TSNR, 9 molecules were analyzed across multiple frames. Error bars for SSNR indicate the standard error, while those for TSNR represent the standard deviation.

As a result, smaller molecules disperse their fluorescence signals across more pixels, while larger molecules focus relatively stronger signals in fewer pixels, which leads to a higher background standard deviation spread (Fig. 2B). We confirmed that both SSNR and TSNR decreased as the hydrodynamic radius increased (Fig.2C and D). When we plotted SSNR against concentration for each hydrodynamic radius, we observed that the decline in SSNR with increasing concentration was more significant for larger molecules. This suggests that studies involving larger molecules encounter greater challenges when using high concentration. Additionally, the larger differences in TSNR at low concentrations indicate that time-series intensity analysis becomes more difficult for larger molecules, even at low concentrations.

### Improved SNR by enhanced molecular motion

Based on our observation that diffusion impacts SNR, we hypothesized that increasing molecular motion could improve SNR. To test this, we introduced linear velocity along the x-axis to stimulate the diffusion of 20 nM of 1.3 kbp dsDNA in a fluid stream (Fig. 3A). The diffusion coefficient of 1.3 kbp dsDNA was obtained from a previous study [15]. The simulation of fluid flow on background molecules resulted in a substantial increase in both SSNR and TSNR (Fig. 3B, C). At a flow rate of 1 ml/min for 20 nM 1.3 kbp dsDNA, the SNR was comparable to that at a concentration of 1 nM without flow, indicating that accelerating the movement of background molecules can enhance SNR.

**Fig. 3.**
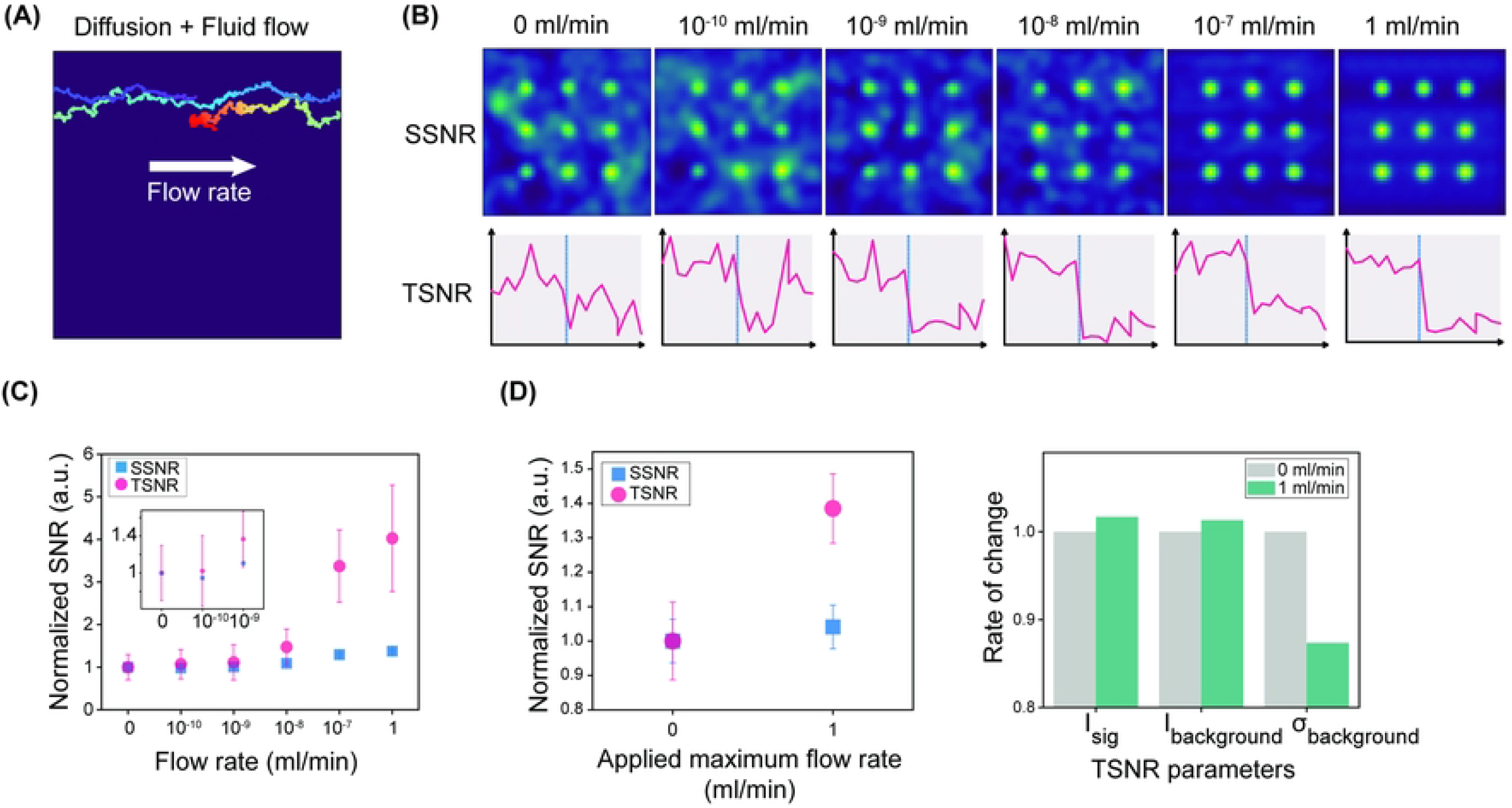
SNR Improved by faster molecular movement. (A) Representative trajectory of a diffusing molecule in the simulation with a fluid flow rate of 10^−7^ ml/min. The diffusion coefficient was calculated using the hydrodynamic radius of 1.3 kbp dsDNA. (B) Representative output images of 20 nM 1.3 kbp DNA in different flow rates. The vertical blue line on the TSNR plot distinguishes signal on and off frames. (C) Normalized SSNR and TSNR in various flow rate simulations. Inset is the magnification of low range of flow rate. SSNR was calculated same as Fig. 1 in 10 frames of output images. TSNR simulation was 20 frames long. In the first half of the image sequence, signal data was added while in the second half there were only background molecules. The first 10 frames were considered as signal on frames, and the last 10 frames as off frames. Error bar of SSNR is standard error and error bar of TSNR is standard deviation. (D) (Left) Experimental SSNR and TSNR of 20 nM 1.3 kbp dsDNA. The applied maximum flow rate was the value of syringe pump. For SSNR, 391 frames were calculated and averaged for 38 molecules. For TSNR, 34 molecules of 0 ml/min, 32 molecules of 1 ml/min were analyzed. (Right) TSNR parameters.

To apply this experimentally, we introduced a fluid flow of 1 ml/min into the imaging chamber. This resulted in a 38.5% improvement in TSNR and a 4.1% improvement in SSNR, attributed to reduced background intensity fluctuations (Fig. 3D). These enhancements correspond to a stimulated velocity between 10^−8^ and 10^−9^ ml/min. Given the parallel plate structure of our flow chamber, the velocity profile of the fluid was described by the equation: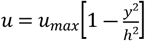, where *h* is half the distance from the center of the chamber to the end of the z-axis, and *y* is the distance from the center along the z-axis[16]. At the imaging plane, the velocity was about 10^―4^ of the center’s velocity. We infer that the decreased SNR enhancement compared to the simulation is due to the lower actual velocity at the imaging plane. Nevertheless, the enhancement of TSNR from the flow effect was on the order of 10 nM, suggesting that further optimization of strategies to accelerate molecular motion could lead to even greater SNR improvements.

## Discussion

Our experiments and computational simulations demonstrate that the diffusion of molecules in solution significantly affects the SNR in single-molecule imaging. The hydrodynamic radius is a key factor influencing the diffusion coefficient, yet it is often overlooked in the design of single-molecule in vitro experiments. Based on our findings, we suggest that immobilizing larger molecules on a surface may yield improved SNR when designing experiments aimed at detecting interactions between two species of molecules.

Conventional wisdom holds that single-molecule imaging becomes challenging at concentrations in the tens of nanomolar range. However, there are examples where high concentration molecules have been successfully imaged. Tokunaga et al. [17] detected ATP-Cy3 in a 50 nM solution, and Lu et al. [18] observed a single ubiquitin-AF647 in a 3 μM solution, though with a relatively long exposure time 3 seconds. These molecules have smaller hydrodynamic radii compared to other biomolecules, which likely explains why sufficient SNR was achieved even at such high concentrations.

In our study, we applied linear flow to the background molecules to investigate the relationship between molecular motion and SNR. While this approach had a limited effect on SSNR, it significantly improved TSNR. For single-molecule kinetics studies, even in cases of poor SNR, methods such as the walking average technique, as used in this study, can help pinpoint the positions of halted molecules across image stacks.

However, resolving molecular binding events, which are essential for kinetic measurements, remains more challenging. Our results indicate that accelerating molecular motion can enhance TSNR, providing a potential method for improving kinetic analysis in these experiments.

Since laminar flow is less effective in accelerating molecular movement and required a large sample volume, enhancing diffusion can be a more efficient way for improving SNR. For instance, introducing active particles into a solution can increase diffusion [19, 20], thereby enhancing SNR. We anticipate that further development of efficient diffusion-enhancement techniques could lead to even greater improvements in SNR for single-molecule imaging.

## Acknowledgements

This research was supported by the National Research Foundation of Korea funded by the Ministry of Science and ICT (J.B.L.: RS-2023-00218927 and J.H.J. RS-2023-00218927 and RS-2024-00343900).

## Author contributions

G.B. and G.K. performed all experiments and computational simulations. S.P. under the J.H.J. guidance provided the simulation code. G.B., G.K, J.-B.L. analyzed data and wrote the paper.

